# Growing Phylogenetic Trees Using Hierarchical Clustering

**DOI:** 10.1101/2022.02.08.479565

**Authors:** Onno Eberhard

## Abstract

We compare mitochondrial DNA of different species to build a phylogenetic tree. The main challenge is that the calculation of the Levenshtein distance is very slow for large sequences. We introduce an approximate scheme which uses Levenshtein for lower-level details and a fast distance based on sequence length for high-level attributes. The tree is built using average linkage hierarchical clustering.

## 1 Introduction

Evolution is a fascinating topic. Darwin’s seminal treatise on the topic [1] is “surprisingly” recent, given that the question of where life and its diversity comes from must be an ancient one. At Darwin’s time, all evidence of evolution was external: slight variations across species hinted at relatedness. Now, with bioinformatics being a completely independent research field, much has changed. We shall exploit two big such changes, which Darwin was not able to do at the time:

1. The availability of genome data. Raw genome data are strings consisting of the letters `A, C, G` and `T`. This data represents the base pairs in an organism’s DNA, and mostly encodes information about the structure of proteins.
2. The availability of computers. We will use computers to recognize patterns in the data and automatically determine the relatedness of species by comparing their DNA sequences.

The goal of this work is to build a phylogenetic tree: a “tree of life”. To figure out the relatedness of two organisms, we want compare their genomes and calculate a relatedness score. In Section 3, we discuss how this score should be calculated, and how it can be used to build a tree.

## 2 Datasets

One problem with DNA data is that it does not naturally fit into a tabular arrangement. DNA is typically distributed across chromosomes. Different species have different numbers of chromosomes, and some species don’t even have DNA, but RNA. This diversity is a result of the diversity of life, but we cannot focus on everything here. To simplify the problem, we restrict ourselves to eukaryotes. This class of species contains all plants, fungi and animals. One major advantage of of this simplification is the availability of mitochondria, which are part of the eukaryotic cell, and are responsible for converting sugar into ATP (the molecule used for energy exchange within a cell). An interesting property of mitochondria is that they possess their own DNA, separate from the other chromosomes in the cell. From here on out, we will restrict ourselves to analysing only mitochondrial DNA.

The genome data we use is provided by the National Center for Biotechnology Information (NCBI) [2]. We provide a script to download and clean the data online^1^. All scripts are explained on the GitHub page. Later, we will also use taxonomy data from the NCBI.

## 3 Method

How do we figure out how related two species are given their DNA sequences? Comparing two DNA strings is usually a very complicated task, with need for sophisticated sequence alignment algorithms. However, as we are only analyzing mitochondrial DNA, the complexity reduces greatly and we may assume a simplistic model of evolution. Under this model, we only consider point mutations and ignore other factors like sexual reproduction (mitochondria reproduce asexually), and copying/deleting of whole chunks of DNA. A point mutation is a “single-character edit”, i.e. inserting, deleting or changing a single character of DNA. The Levenshtein distance *d*_Lev_(*a, b*) is defined as the minimum number of single-character edits to change the string a into b. If we calculate this distance between each pair of DNA sequences in our dataset, we have a reasonable relatedness score under our evolution model. Given such a distance measure, a phylogenetic tree can be built using (agglomerative) hierarchical clustering (Algorithm 1). We use average-linkage clustering which means that the distance of two clusters is the average of the distances of species in this cluster. This is a natural choice under our evolution model.

### Algorithm 1: Hierarchical Clustering

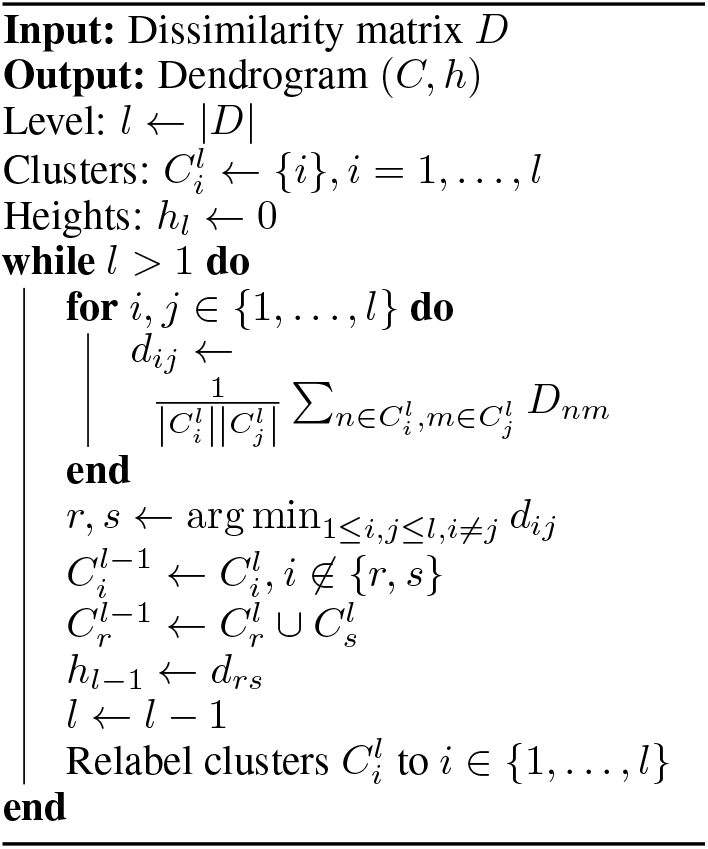

One big problem with using the Levenshtein distance is that it is slow to compute, even when using fast C libraries. Especially for long sequences (≥ 40.000) it becomes prohibitive. To solve this problem, we try to approximate the Levenshtein distance by a very simple heuristic, which we call the length-distance: *d*_len_(*a, b*):= |len(*a*) – len(*b*)|. The rationale behind this is that for large sequence differences, the Levenshtein distance will be dominated by the difference in length. Both distance measures are compared to the real taxonomy classification in Figure 1 on a small test set.

**Figure 1:**
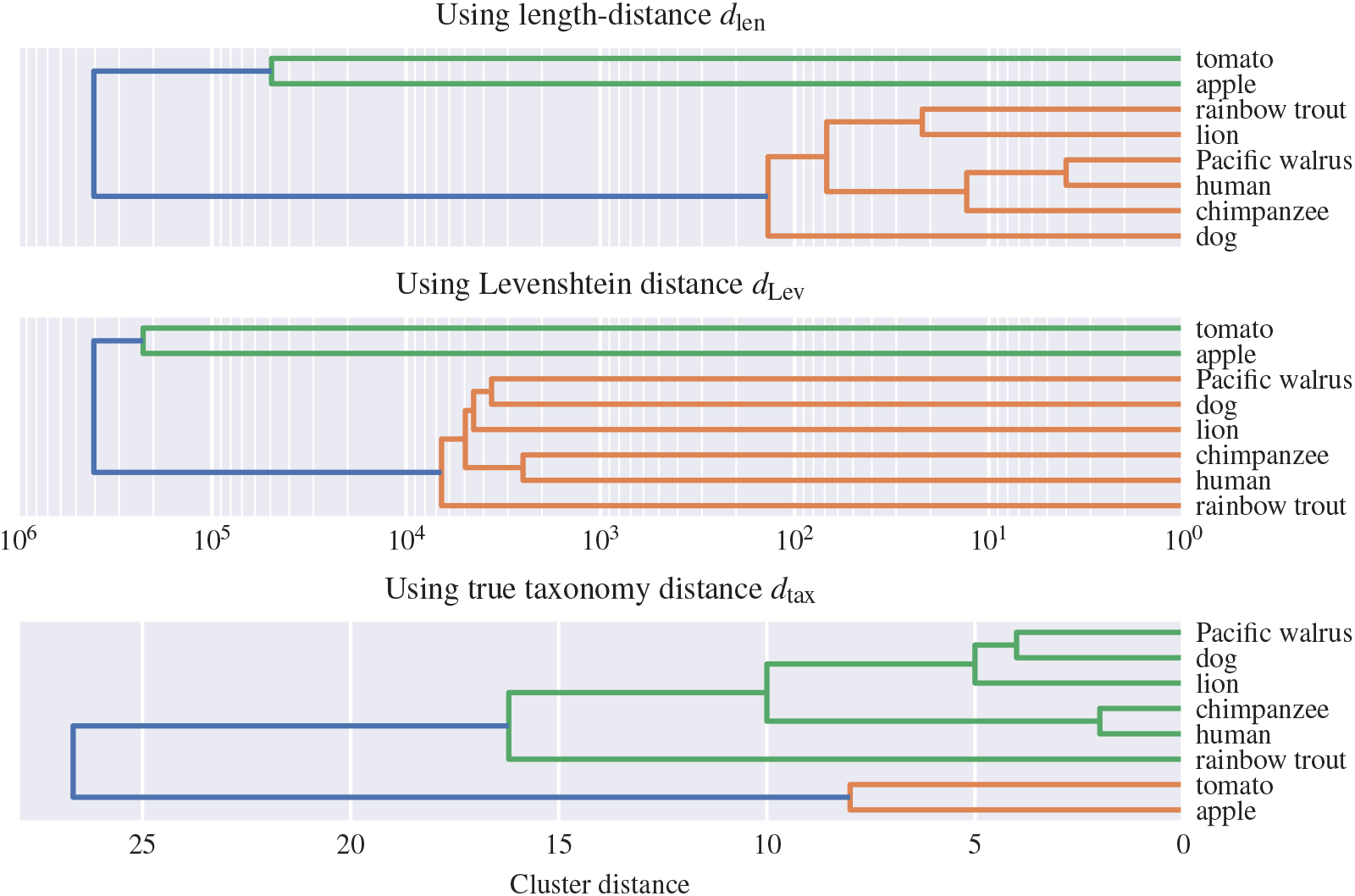
The length-distance works well on the macro- but not on the micro-level.

As the length-distance seems to work okay on large-scale structure, we ideally would want to only use the Levenshtein distance on shorter sequences. To this end, we analyze the distributions of length-distances in the left plot in Figure 2. A natural cut-off point is to use Levenshtein for all pairs *a, b* where *d*_len_ < 2000. We will also filter out pairs where 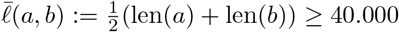, as shown in the center plot (Fig. 2). In the right are shown the Levenshtein distances (after 24h of computation), where we filter the lonely outliers of *d*_Lev_ ≥ 20.000. Finally, we define:

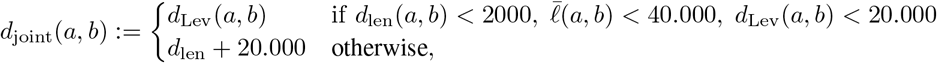

which is the joint distance of *d*_len_ and *d*_Lev_. It is clear that always *d*_Lev_ ≥ *d*_len_, as the Levenshtein distance must at least account for the difference in length. Thus, we add 20.000 to the length-distance, so as not to interfere with the smaller scale. A comparison with *d*_tax_, which we define as the number of true taxonomy classes that *a* and *b don’t* have in common, is shown in Figure 3.

**Figure 2:**
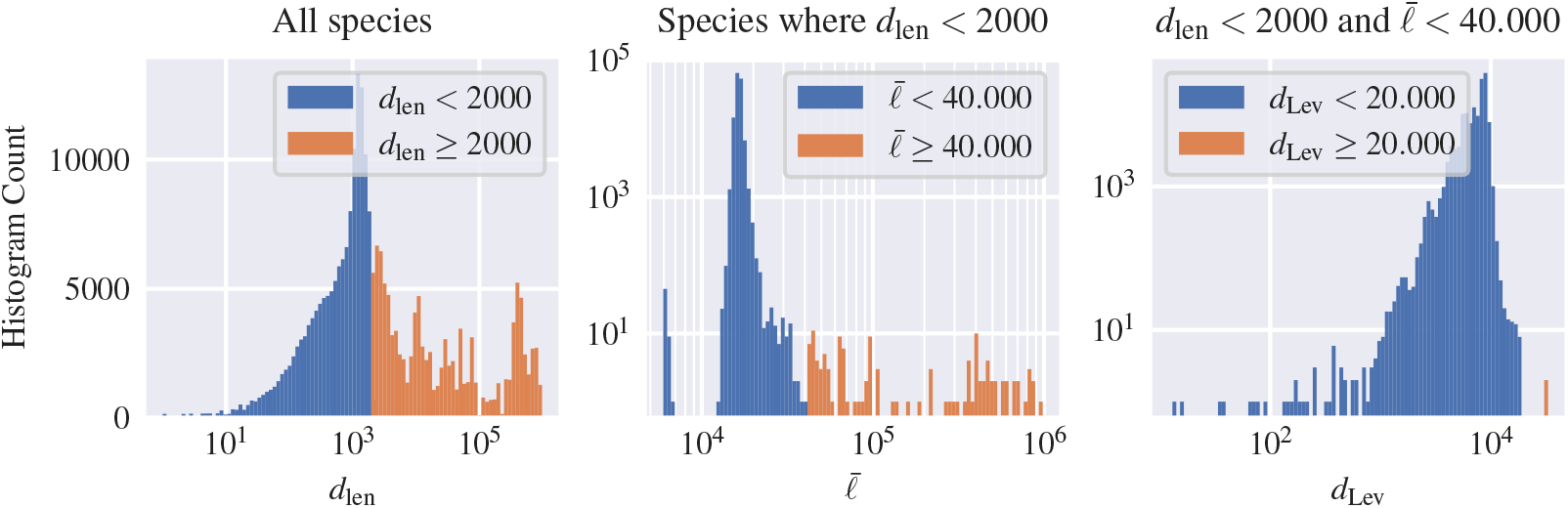
We must decide where to trade off accuracy (*d*_Lev_) for efficiency (*d*_len_).

**Figure 3:**
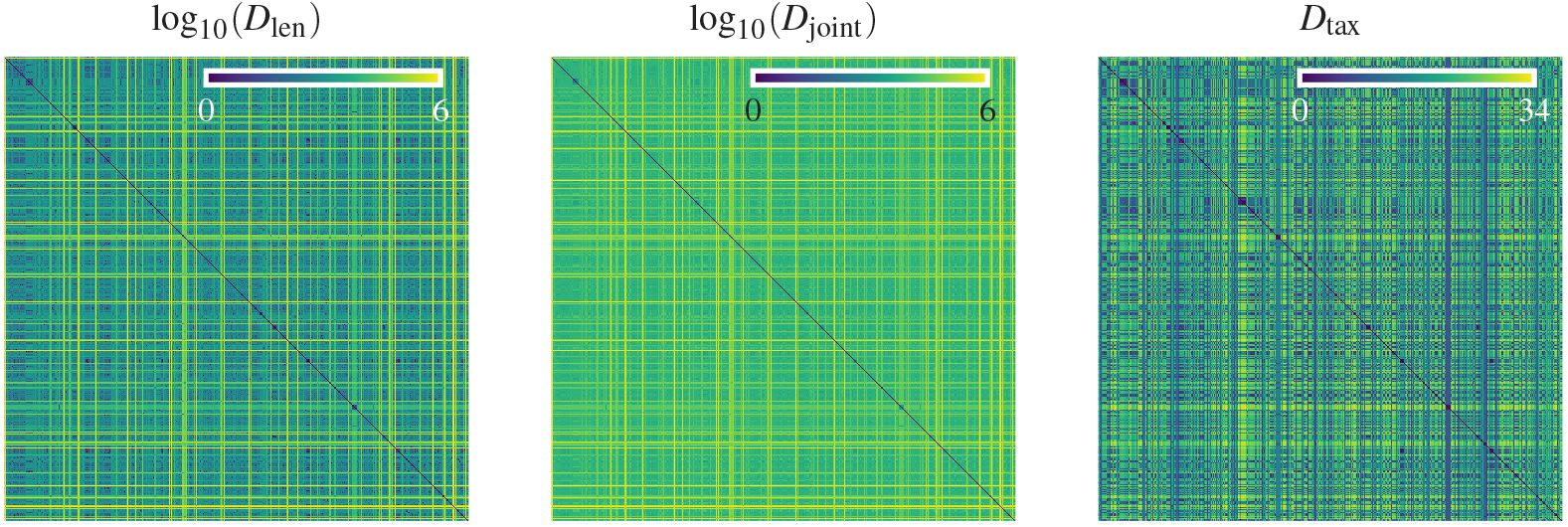
Distance matrices created for the species dataset 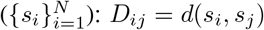.

## 4 Results

The resulting phylogenetic tree is shown in Figure 4, where it is compared to one built with only the length-distance. It can be seen that using the *d*_joint_ works in practise, where calculating the Levenshtein distance is not feasible. However, there might be other computationally efficient distance measures that perform better than *d*_len_. Our evolution model is also not perfect, as DNA mutations are not always point mutations, but chunks can be copied or moved as one. A better distance measure would respect this as valid *single* mutations.

**Figure 4:**
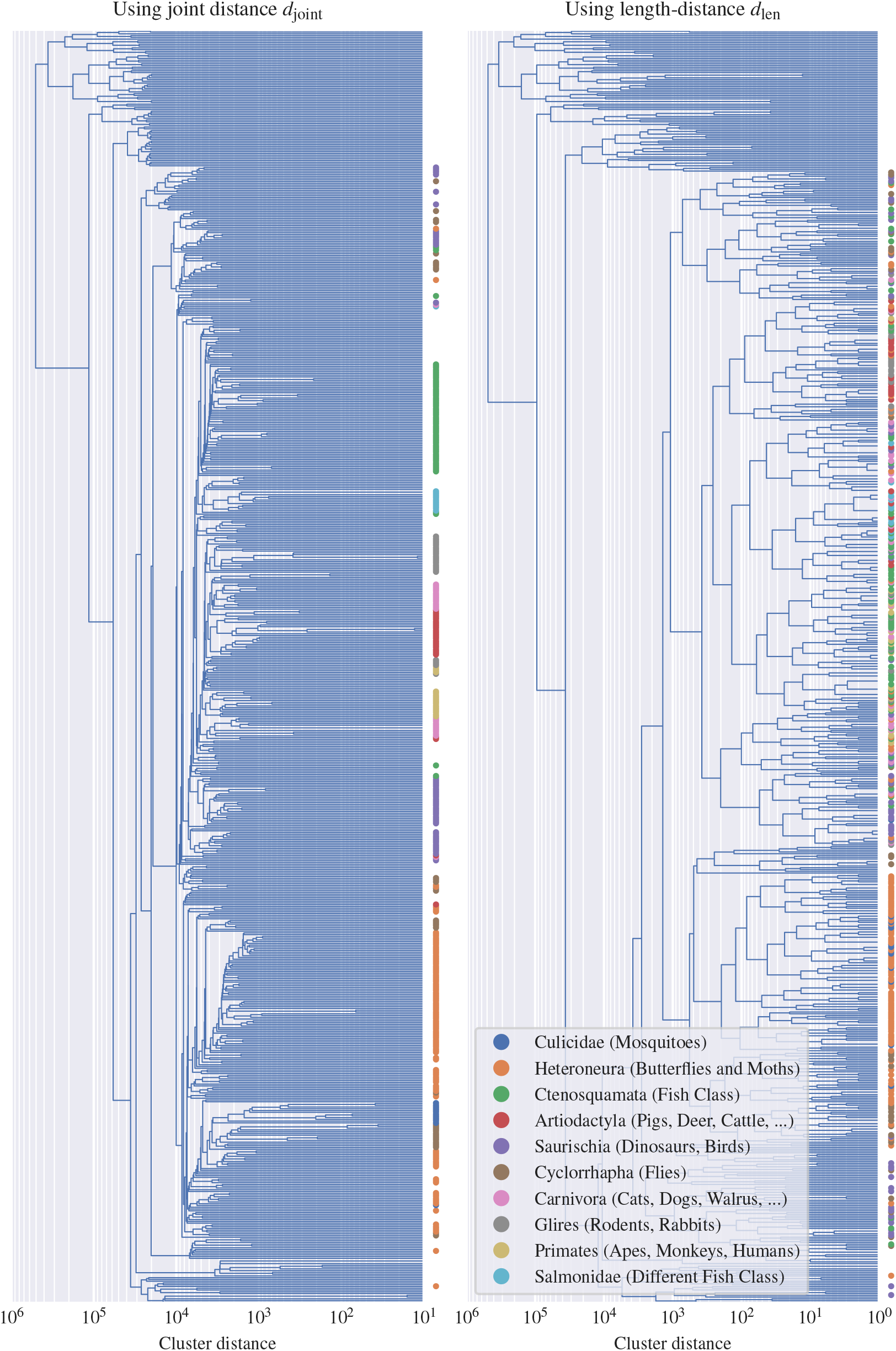
Phylogenetic trees built using hierarchical clustering. Plants and fungi, which are at the top of the plots, are correctly separated from animals in both cases. The colored dots represent known taxonomy groups and should ideally be close together (even form convex sets) in the final tree. As expected, *D*_joint_ works much better than *D*_lev_, though some dots are still distributed a bit, which might show that even for higher-level structure, the length-distance is suboptimal.

## Footnotes

1 https://github.com/onnoeberhard/phylo

